# The food web of Potter Cove (Antarctica): complexity, structure and function

**DOI:** 10.1101/094557

**Authors:** Tomás I. Marina, Vanesa Salinas, Georgina Cordone, Gabriela Campana, María Eugenia Moreira, Dolores Deregibus, Luciana Torre, Ricardo Sahade, Marcos Tatián, Esteban Barrera Oro, Marleen De Troch, Santiago Doyle, María Liliana Quartino, Leonardo A. Saravia, Fernando R. Momo

## Abstract

Knowledge of the food web structure and complexity are central to better understand ecosystem functioning. A food-web approach includes both species and energy flows among them, providing a natural framework for characterizing species’ ecological roles and the mechanisms through which biodiversity influences ecosystem dynamics. Here we present for the first time a high-resolution food web for a marine ecosystem at Potter Cove (northern Antarctic Peninsula). Eleven food web properties were analyzed in order to document network complexity, structure and topology. We found a low linkage density (3.4), connectance (0.04) and omnivory percentage (45), as well as a short path length (1.8) and a low clustering coefficient (0.08). Furthermore, relating the structure of the food web to its dynamics, an exponential degree distribution (in- and out-links) was found. This suggests that the Potter Cove food web may be vulnerable if the most connected species became locally extinct. For two of the three more connected functional groups, competition overlap graphs imply high trophic interaction between demersal fish and niche specialization according to feeding strategies in amphipods. On the other hand, the prey overlap graph shows also that multiple energy pathways of carbon flux exist across benthic and pelagic habitats in the Potter Cove ecosystem. Although alternative food sources might add robustness to the web, network properties (low linkage density, connectance and omnivory) suggest fragility and potential trophic cascade effects.

## 1. Introduction

Food web (FW) characterization is essential to understanding ecology as a way to describe and quantify the complexity of ecosystems by identifying the trophic interactions among species (Bascompte 2009). The framework of ecological network analysis could also be used to quantify the effects of the environment and how indirect effects of such interactions influence overall ecosystem properties (Brose and Dunne 2009).

Since the early 2000s, ecological networks from marine systems have received more attention answering an emphatical call of Raffaelli (2000) for more research on marine webs. In this sense, indices derived from Ecological Network Analysis (ENA), a system-oriented methodology to analyze within system interactions (Fath et al. 2007), have been used to investigate trophic interactions in marine ecosystems (Baird et al. 2007, Ulanowicz 2011, Wuff et al. 2012, Heymans et al. 2014). Among marine webs, polar FWs recently began to be considered in the frame of FW theory (e.g. Jacob et al. 2006, Bodini et al. 2009, de Santana et al. 2013). Moreover, some conclusions on the effects of global warming on Arctic and Antarctic marine FWs have been proposed (de Santana et al. 2013, Kortsch et al. 2015).

Potter Cove is an Antarctic fjord that suffers from the impact of the high rate of warming occurring in Western Antarctic Peninsula (Quartino et al. 2013, Deregibus et al. 2016). The abundant and rich epibenthic fauna has been changing under the influence of considerable sediment inputs and other effects derived from ice melting (Pasotti et al. 2015a, Sahade et al. 2015). The way in which network properties can be modified under climate change is in general, poorly known (Petchey et al. 2010, Walther 2010, Woodward et al. 2010). To understand the community-level consequences of the rapid polar warming, Wirta et al. (2015) suggested that we should turn from analyses of populations, population pairs, and isolated predator–prey couplings to considering all the species interacting within communities. If species affected by perturbations possess key functional roles in the FW, then the potential higher order, indirect effects of those perturbations on the entire FW structure can be dramatic (Kortsch et al. 2015). Knowing that climate change effects are already occurring in Potter Cove ecosystem and that ecosystems respond to perturbations as a whole, a network approach could contribute to a better understanding of changes in the ecosystem’s synthetic properties like resilience or stability. A representative roadmap of trophic interactions of Potter Cove will allow testing for the impact of ongoing climate change effects (e.g. glacier retreat, loss of ice shelves, increment of sedimentation input) which might be transmitted throughout the entire ecosystem.

Although FW studies use binary webs that indicate the presence of a trophic interaction but do not provide any information on the frequency of the interaction or the rate of biomass flow through the interaction, overlap graphs (e.g. competition and common-enemy graphs), can provide information about indirect interaction strength between predators and prey, respectively. Indirect effects in predator and prey assemblages can also be studied by evaluating these graphs. The strength of predator-predator and prey-prey indirect interactions is extremely difficult to measure but, if they prove generally prevalent, they could be a major driver of community dynamics and ecosystem functioning (Woodward et al. 2005). The analysis of the degree distribution of links in the overlap graphs, omitted in most FW studies, might be very useful to identify, based on the competition graph, generalist and specialist predators, and to evaluate energy pathways in the common-enemy graph.

In the current work, we present the first, detailed analysis of the FW for the Potter Cove ecosystem (South Shetland Islands, Antarctica). The objectives of this study were to: 1) analyze the complexity and structure of the ecological network in the context of the most-studied marine FWs; and 2) examine its degree distribution and overlap graphs in order to gain insight into the ecosystem dynamics and functioning.

## 2. Methods

Potter Cove is a 4 km long and 2.5 km wide Antarctic fjord located at 25 de Mayo/King George Island (62°14´S, 58°40´W, South Shetland Islands) (Fig. 1). A shallow sill (< 30 m) separates its inner and outer areas. The inner cove is characterized by soft sediments and by a shallower depth than the outer cove (< 50 m); in the outer cove the bottom is mainly rocky and with average depths of 100 m. Potter Cove is adjacent to Maxwell Bay, which connects to the Bransfield Strait. Water circulation in Potter Cove is strongly influenced by the general circulation of Maxwell Bay (Roese and Drabble 1998). A cyclonic circulation has been identified, with efficient water renewal in the northern sector, where water from Maxwell Bay enters the Cove. Freshwater input varies both seasonally and inter-annually and carries important amounts of suspended sediments. Two main creeks discharge into the Cove, the Matias and the Potter creeks. They exhibit different regimes, the first being snowy and lacustrine, the latter snowy and glacial (Varela 1998). Drainage ranged between 0.03 and 0.11 m^3^ s^−1^ in the Matias Creek and from 0.08 to 3.8 m^3^ s^−1^ in Potter Creek (Varela 1998). Suspended sediment discharges ranged between 0.04 and 15 kg m^−3^ (average = 0.14 kg m^3^), which correlate with air temperature. These characteristics are consistent with data from other glaciomarine environments in Antarctic coastal waters (Leventer and Dunbar 1985).

**Fig. 1.**
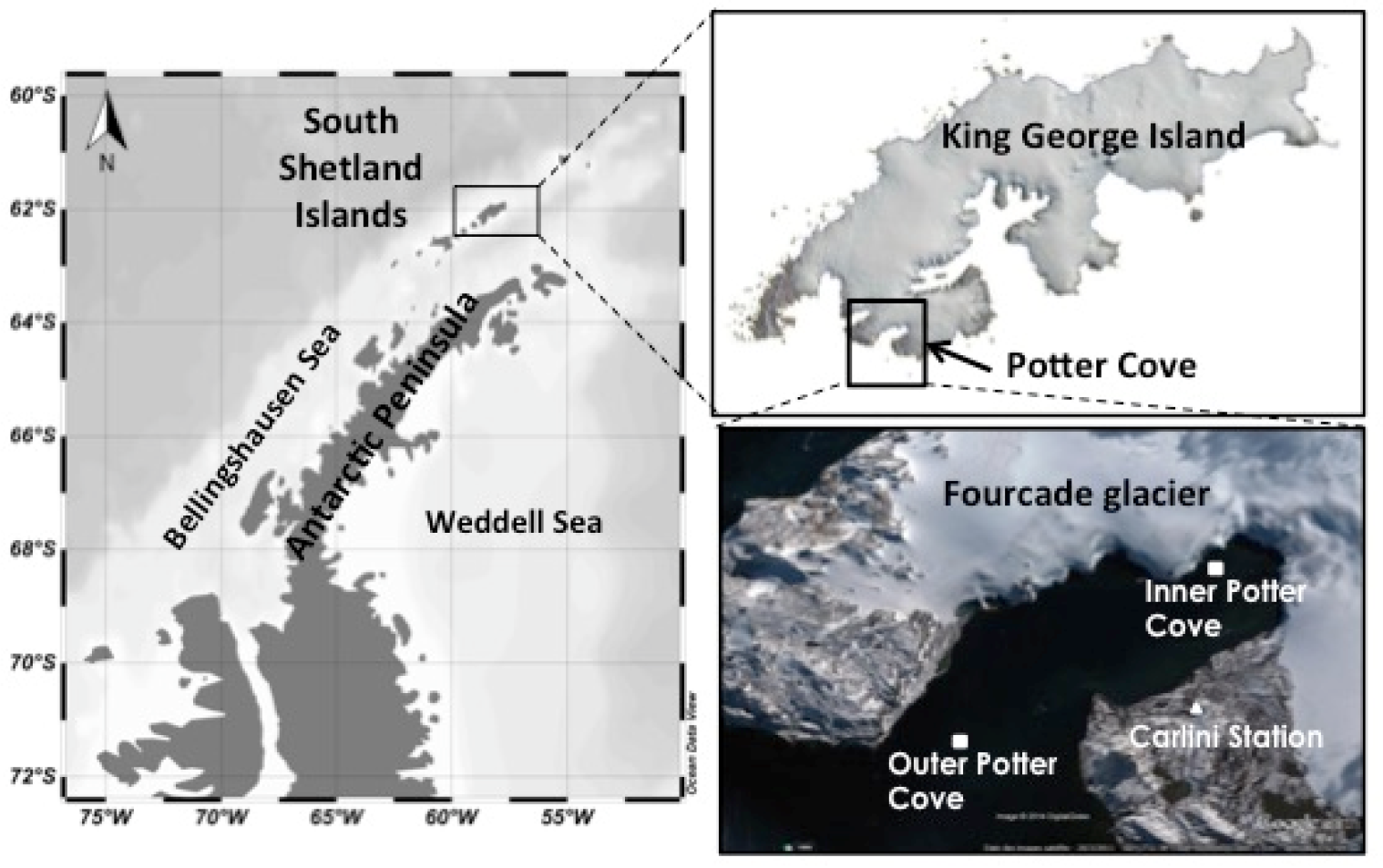
Map of Potter Cove and its location on Isla 25 de Mayo/King George Island.

### 2.1. Food web assembly

We constructed the FW of Potter Cove ecosystem primarily based on information about species living in that system and their feeding habits from studies within the framework of an international research cooperation between Argentina and Germany initiated in 1994 and ongoing for more than 20 years (Wiencke et al. 1998, 2008).

We collected information on feeding links by a thorough literature search (> 500 papers and reports revised). To assemble the network we only considered trophic interactions confirmed by gut content studies and/or field observation. Furthermore, direct observations of researchers from field sampling campaigns in the Cove (e.g. divers when collecting benthic samples) were also taken into account. Laboratory experimental studies, where feeding selectivity, palatability or behavior was tested, were not included in this study as we consider the trophic links proved from experiments are not as robust as the ones gathered from the field data. Investigations using biomarkers (i.e. stable isotopes and fatty acids) were not considered since trophic interactions are established by sampling few individuals (n ≈ 10-100) and studied prey-predator relationships are usually between trophic species widely aggregated. Further details on the trophic links included in the present study (references and methods used to confirm a link) are presented in the electronic supplementary material (Appendix A).

Trophospecies, here defined as aggregated groups of taxa, were only considered when data on specific biological species were not available (lack of data resolution) or when taxa shared the same set of predators and prey within the FW (trophic similarity criteria). We have not considered top vertebrate predators (e.g. penguins, seals, whales), as they only sporadically enter the Cove to feed. In addition, pelagic fish (typically taken by Antarctic penguins and pinnipeds) were not considered due to paucity of ocurrence (Barrera-Oro and Casaux 2008).

The diversity of the expertise of the authors contributing to the present study was a key factor in generating the quality of the FW, and inherently improved the network representation of the Potter Cove ecosystem.

### 2.2. Network analysis

An interaction matrix of pairwise interactions was constructed; a value of 1 or 0 was assigned to each element *a*_*ij*_ of the matrix depending on whether the *j*-species preyed or not on the *i*-species. The FW is an oriented graph with *L* trophic links between *S* nodes or species. The FW graph was drawn from the interaction matrix using Visone software version 2.9.2 (Brandes and Wagner 2004).

Several network properties that are commonly used to describe complexity and structure in FWs were calculated (Dunne et al. 2002b, de Santana et al. 2013): (1) number of species, *S;* (2) total number of interactions or trophic links, *L*; (3) number of interactions per species or linkage density, *L/S*; (4) connectance or trophic links divided by total number of possible interactions, *C*=*L/S^2^*; percentage of (5) top species (species with prey but without predators), (6) intermediate species (species with prey and predators), (7) basal species (species with predators/consumers but without prey); and (8) percentage of omnivores (species eating prey from more than one trophic level).

Trophic levels (*TL*) of species were calculated using the short-weighted *TL* formula of Williams and Martinez (2004). Short-weighted trophic level is defined as the average of the shortest *TL* and prey-averaged *TL*. Shortest *TL* of a consumer in a food web is equal to 1 + the shortest chain length from this consumer to any basal species (Williams and Martinez 2004). Prey averaged *TL* is equal to 1 + the mean *TL* of all consumer´s trophic resources, calculated as

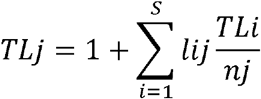

where *TL*_*j*_ is the trophic level of species *j*; S is the total number of species in the food web; *l*_*ij*_ is the connection matrix with *S* rows and *S* columns, in which for column *j* and row *i*, *l*_*ij*_ is 1 if species *j* consumes species *i* and 0 if not; and n_*j*_ is the number of prey species in the diet of species *j*. Therefore, Short-weighted *TL* yields a minimum estimate of *TL* and assumes a value of 1.0 for basal species (Williams and Martinez 2004). We considered the mean *TL* of the web as the average of all species’ *TL*.

Two secondary graphs, the competition graph and the common-enemy graph, were constructed. The first one, also known as predator overlap graph, connects predators that share one or more prey, while the latter is drawn by connecting prey species sharing one or more predators (Pimm et al. 1991). Predator overlap graph results were discussed considering dietary data on each predator species involved. To examine a plausible organization in predator and prey species, we separately studied the degree distribution of links *P(k)* for each overlap graph. Links in predator distribution represent the number of prey, while in prey distribution it depicts number of predators. Graphs were plotted using Visone software (version 2.9.2).

We also studied the topology of the FW by measuring three more properties: (9) characteristic path length (ChPath), or the average shortest path length between all pairs of species, (10) clustering coefficient (CC), or the average fraction of pairs of species connected to the same species that are also connected to each other, and (11) degree distribution, or the fraction of trophic species P(*k*) that have *k* or more trophic links (both predator and prey links) (Albert and Barabási 2002). Trophic links were treated as undirected when calculating path length and clustering because effects can propagate through the web in either direction, through changes to both predator and prey species (Watts and Strogatz 1998).

Results of these properties and the ones aforementioned for Potter Cove FW were compared among other marine webs that were chosen considering different criteria: size (S > 25), temporal era (fourth era, see Link et al. 2005) and quality data (i.e. FWs built upon stable isotopes were excluded).

Degree distributions of the total FW and of the mentioned overlap graphs were examined and fitted using nonlinear regression analysis (Xiao et al. 2011). Model selection was performed by computing the Akaike Information Criterion corrected for small sample size (AICc) (Burnham and Anderson 2002, Xiao et al. 2010). R package *nls* (Nonlinear Least Squares) was used to make power-law and exponential fitting (R Core Team 2016).

## 3. Results

The Potter Cove FW (Fig. 2) includes 91 species, composed of 71 biological species, 17 trophospecies (i.e., merging two or more taxonomic species by trophic similarity) and 3 non-living nodes (i.e. fresh detritus, aged detritus and necromass).

**Fig. 2.**
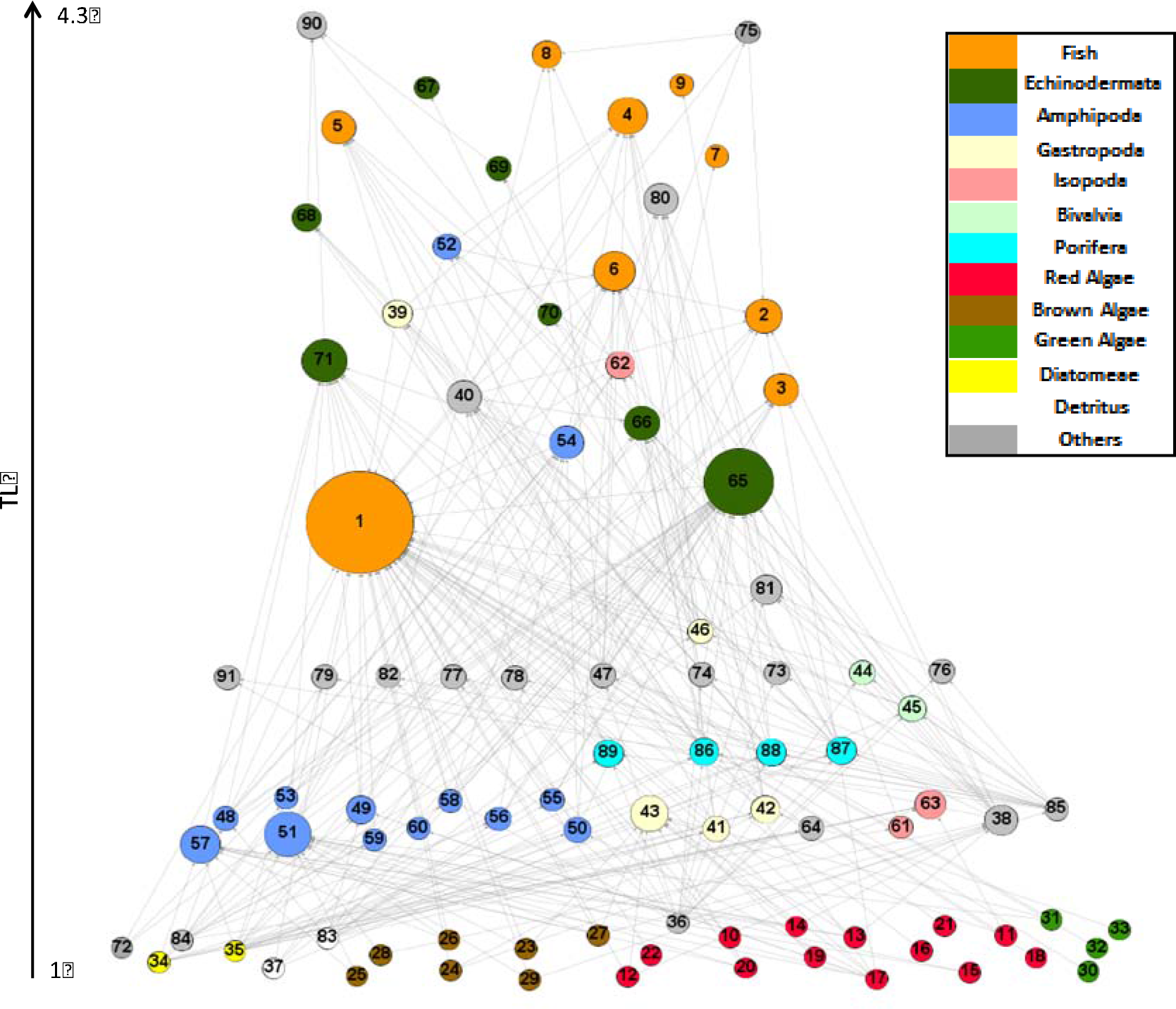
Graphic representation of Potter Cove FW with the trophic level (TL) on the vertical scale and node width proportional to the total degree (in- and out-). Node colors are by functional group. Network was plotted with Visone (version 2.9.2). See electronic supplementary material (Appendix B) for exhaustive lists of trophic species, their trophic level, degree (in- and out-links), functional and taxonomic group affiliation (e.g. algae, phytoplankton, zooplankton, fish, amphipods).

Algae (24 species) comprise red (13 spp.), brown (7 spp.) and green algae (4 spp.). The next trophic levels consist of 13 amphipod species, 3 isopod species, 4 sponge species (one aggregated node: *Stylocordyla borealis* and *Mycale acerata*), 5 gastropod species, 2 bivalve species, 7 echinoderm species, and 9 demersal fish species. See electronic supplementary material (Appendix B) for exhaustive lists of taxa, their trophic level, degree (in- and out-links), functional and taxonomic group affiliation (e.g. algae, phytoplankton, zooplankton, fish, amphipods).

The first thing to note about Potter Cove FW is that most of the species (47%) were at intermediate levels, implying that they act as predators and prey depending on the trophic interaction they are involved in. Moreover, as shown in Fig. 2 some species are far more connected (9 species with degree > 15) than others, according to the total number of trophic interactions they have (e.g. fish and echinoderms).

The main properties of the network complexity for Potter Cove FW included 307 total interactions and a linkage density of 3.4. As a consequence, a connectance of 0.04 was reported (Table 1).

**Table 1.** Properties of network complexity and structure for Potter Cove FW. S = number of trophic species, L/S = linkage density, C = connectance (L/S^2^), T = % top species, I = % intermediate species, B = % basal species, Omn = percentage of omnivorous, TL = mean trophic level, ChPath = characteristic path length, CC = clustering coefficient.

| Food web | S | L/S | C | T | I | B | Omn | TL | ChPath | CC |
| --- | --- | --- | --- | --- | --- | --- | --- | --- | --- | --- |
| Potter Cove | 91 | 3.4 | 0.04 | 19 | 47 | 34 | 45 | 2.1 | 1.8 | 0.08 |

Although intermediate species outnumbered top and basal species, comprising more than half of the species in the FW, the basal species were also numerous (Table 1). In addition, almost half of the species were omnivorous (45%), similar to the percentage observed in intermediate species. The mean trophic level (TL) for Potter Cove FW was 2.1, which was supported by the relatively high proportion of basal species that tend to lower the average.

Network topological properties, characteristic path length (ChPath) and clustering coefficient (CC) were 1.8 and 0.08, respectively.

The degree distribution for the Potter Cove FW (Fig. 3) showed that the exponential model best fitted the data, according to nonlinear regression and AICc analyses (Table 2). The three species with the highest degree were: *Notothenia coriiceps* (fish, 48 links), *Ophionotus victoriae* (echinoderm, 33 links) and *Gondogeneia antarctica* (amphipod, 20 links).

**Fig. 3.**
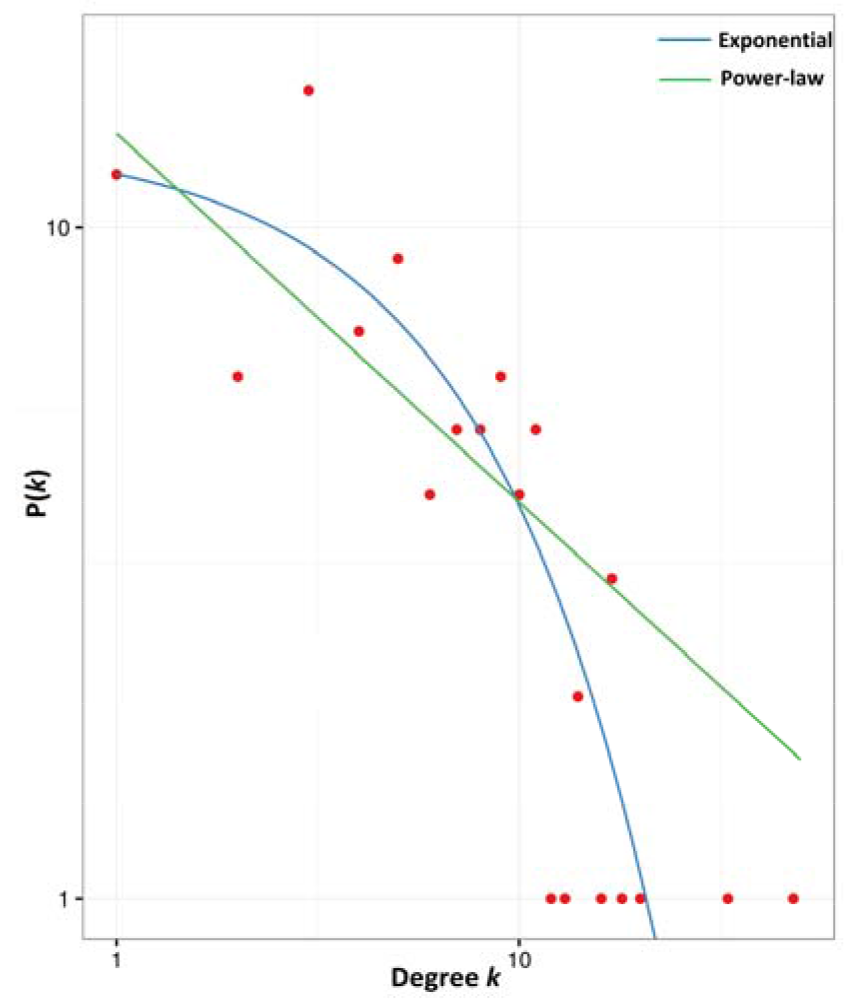
Log-log degree distribution of links *P(k)* for Potter Cove FW. Two candidate models are shown. Best fit is the exponential model.

**Table 2.**
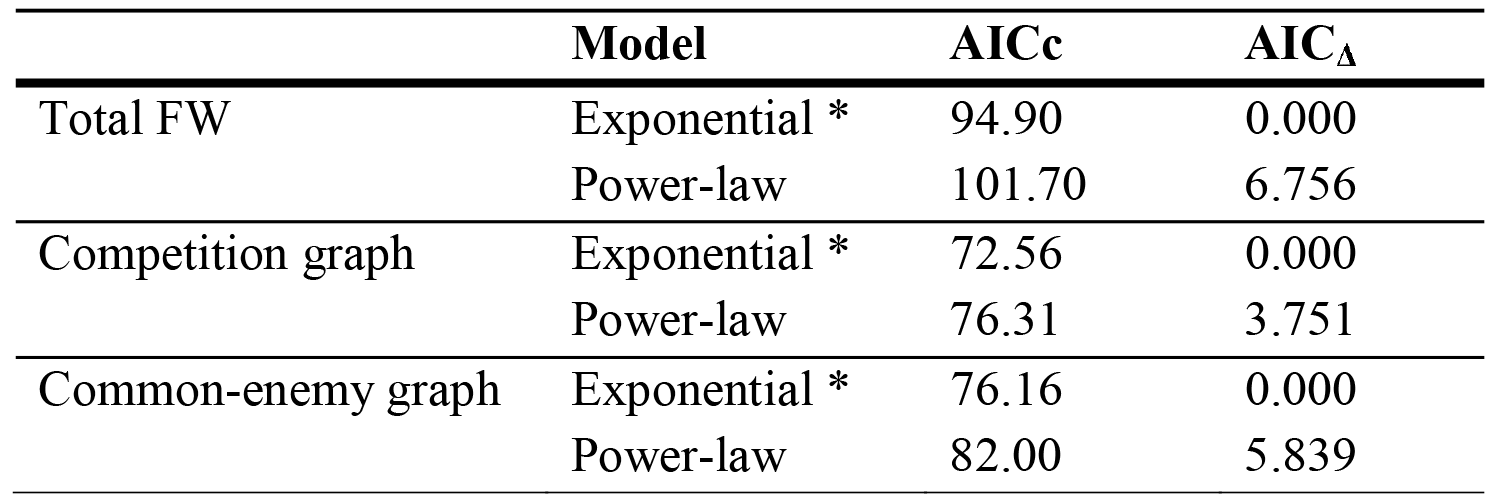
Model fit of exponential and power-law models for degree distributions of total FW (in- and out-links), competition (only predators) and common-enemy (only prey) overlap graphs. AICc and AIC_Δ_ are the Akaike corrected for small sample size and delta values for each candidate model. * Indicates best-fit model.

The competition graph derived from Potter Cove FW is highly connected. It includes 60 species and 478 indirect interactions (Fig. 4) and shows that several pairs of predators share many prey. For instance, all trophic species of sponges form a more connected group than with the rest of the prey species. Furthermore, some species of echinoderms, amphipods and demersal fish are intensively competing for common food sources (see link width and color, Fig. 4).

**Fig. 4.**
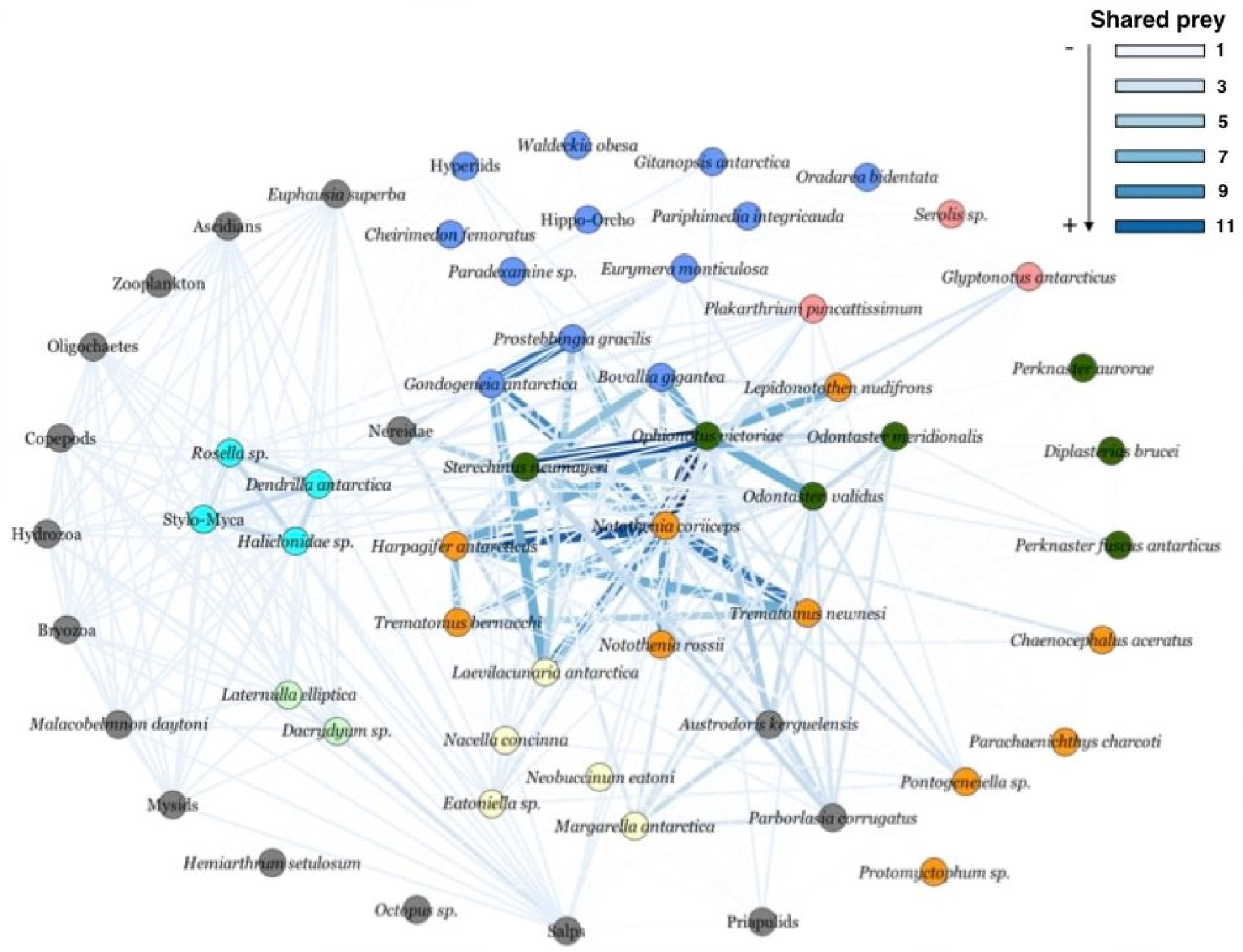
Competition graph for the Potter Cove FW. Node colors (as in Fig. 2): functional groups. Link width and colors: number of shared prey.

To study these potential species interactions, specific competition graphs for the latter two functional groups were built (Fig. 5). The fish overlap graph includes 9 biological species and 28 competitive interactions. It is worthy to note that two species, *Notothenia coriiceps* and *Harpagifer antarcticus*, presented highly overlapping diets. Moreover, *N. coriiceps shares many of the same prey species, which may or may not involve any competition* (Fig. 5a). On the other hand, the amphipod overlap graph suggested low resource overlap among species. However, *Gondogeneia antarctica* and *Prostebbingia gracilis* have many prey in common (Fig. 5 b).

**Fig. 5.**
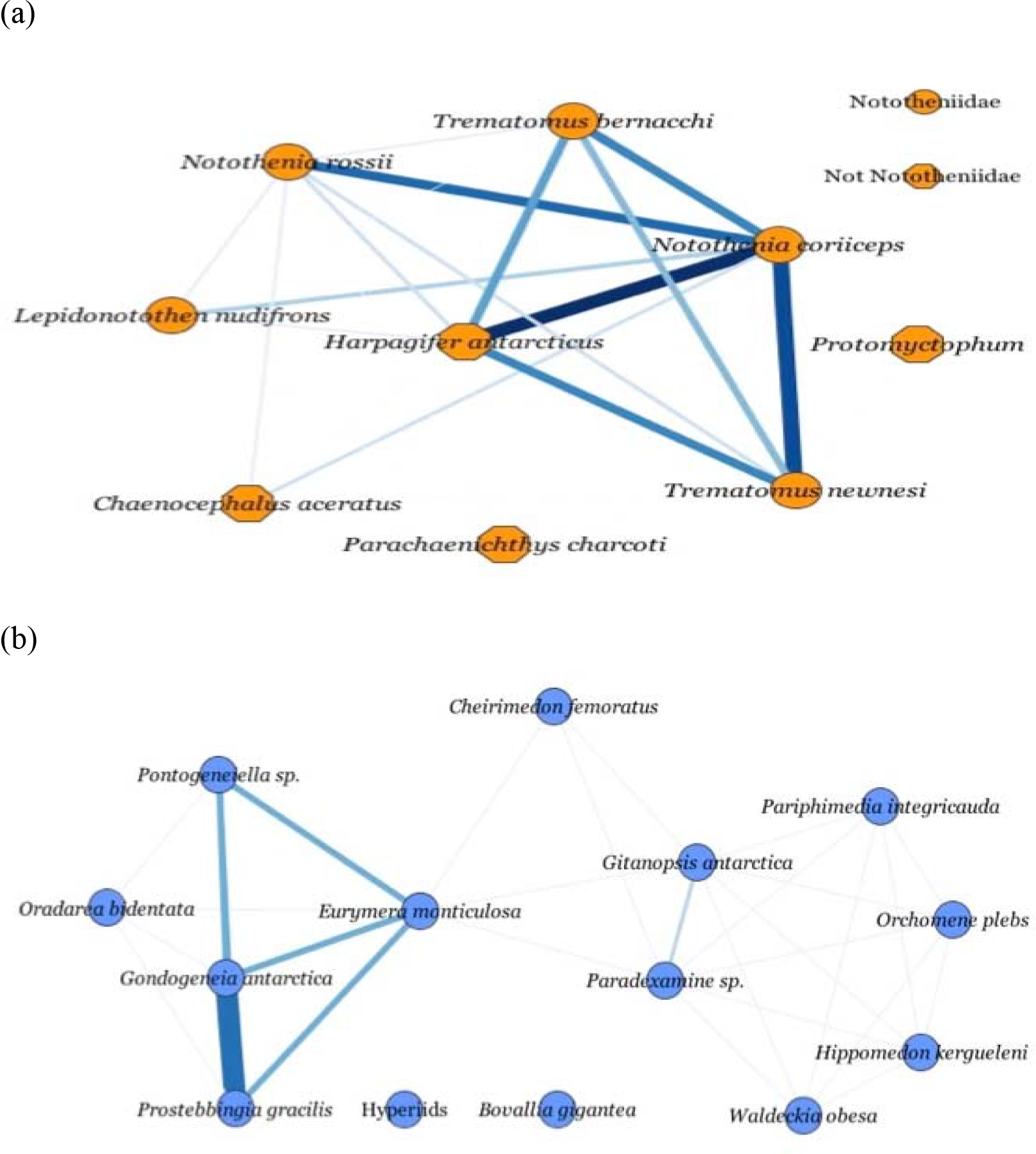
Competition graphs for (a) demersal fish and (b) amphipod functional groups. Link width and colors: number of shared prey (see Fig. 4).

The common-enemy graph shows a hyperconnected structure, where the majority of the species are connected. It contained 74 prey species and 1497 indirect interactions (Fig. 6, up-left). Most of the species are connected due to having only one predator in common. In order to elucidate groups of species having stronger indirect interactions, we eliminated links with value 1. This new graph (Fig. 6, large network) showed groups of species connected by strong interactions: sponges (except for *Dendrilla antarctica*), benthic diatoms – fresh detritus, benthic diatoms – epiphytic diatoms, zooplankton – phytoplankton, some species of amphipods (i.e. *Gondogeneia antarctica* – *Paradexamine sp*. – *Prostebbingia sp*. – *Eurymera meticulosa*), and several red and brown algae (*Gigartina skottsbergii* – *Desmarestia menziesii* – *Iridaea cordata*) (Fig. 6).

**Fig. 6.**
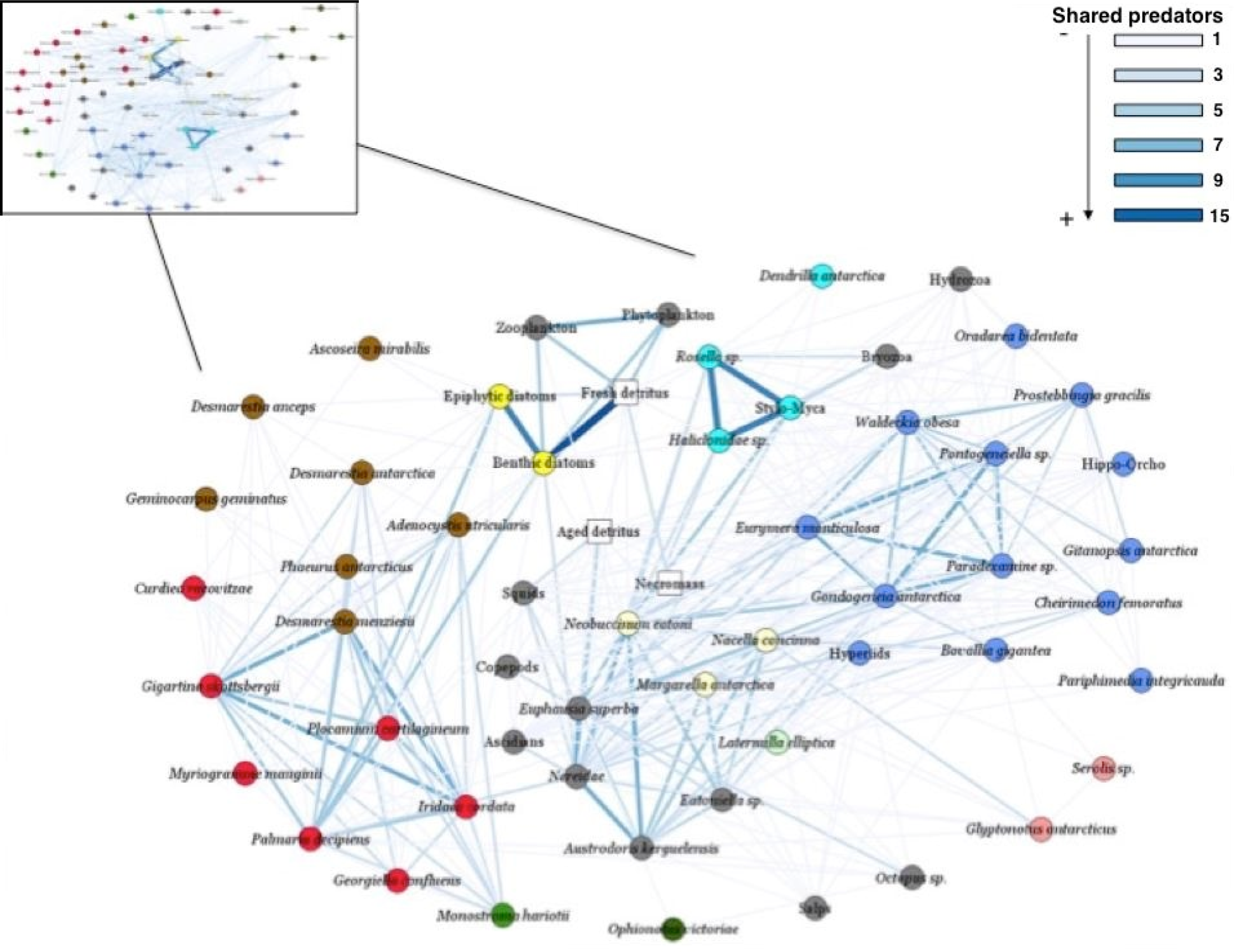
Common-enemy graph for Potter Cove FW. Original graph in left upper corner. Large network shows prey species that share more than one predator. Node colors (as in Fig. 2): functional groups. Link width and colors: number of shared predators.

Degree distribution of links in the competition and common-enemy graphs (Fig. 7) fit best to an exponential model (Table 2).

**Fig. 7.**
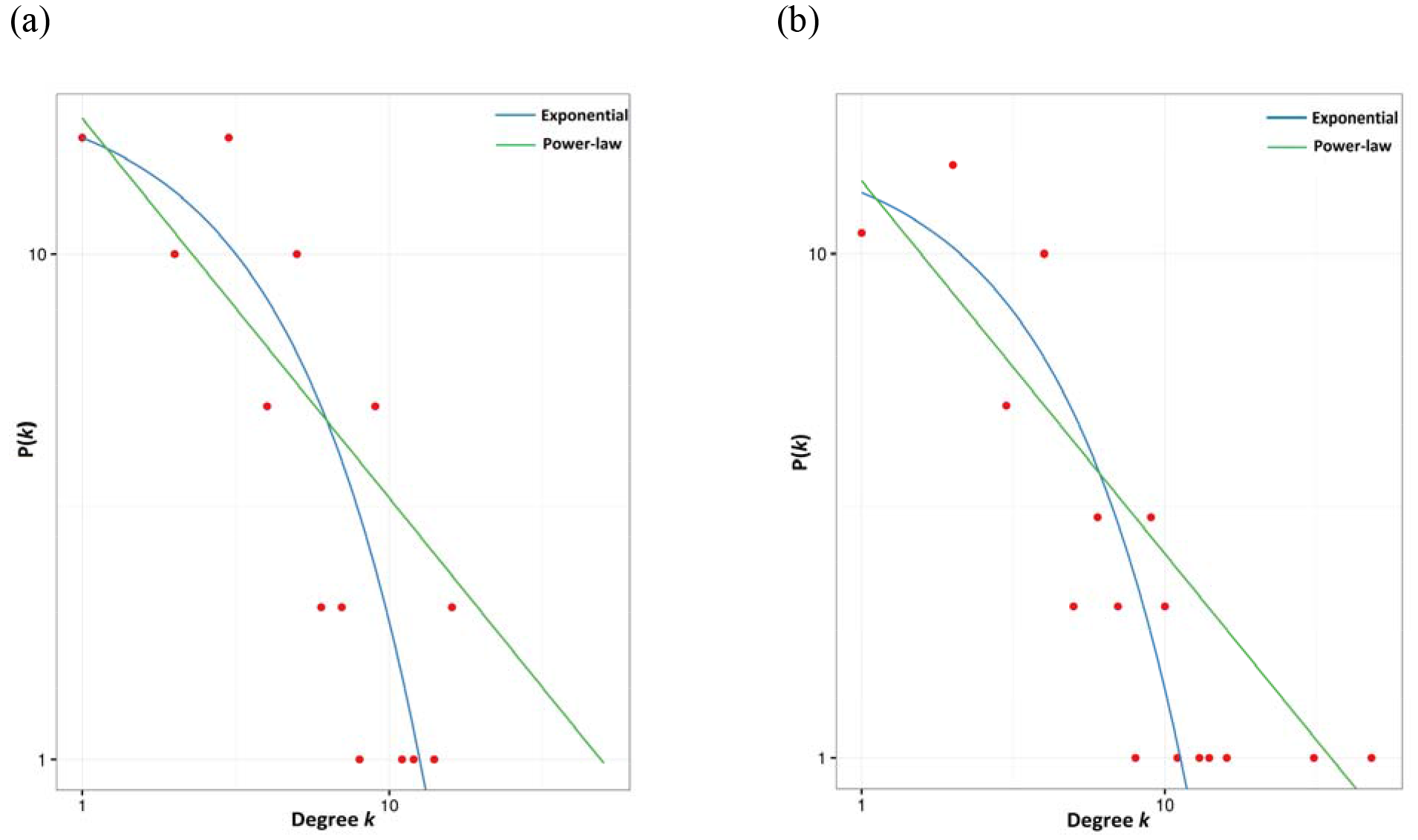
Log-log degree distribution of links *P(k)* for (a) the competition and (b) common-enemy graphs. Best fit is the exponential model for both distributions.

Comparison between the Potter Cove FW and other marine webs showed that linkage density (*L/S*) and connectance (*C*) were lower in the Potter Cove web. The proportions of top and basal species were relatively high, whereas the percentage of omnivory was the second lowest among all webs that were compared. While the characteristic path length in Potter Cove FW was similar to the rest of the FWs, the clustering coefficient was one order of magnitude lower (Table 3).

**Table 3.**
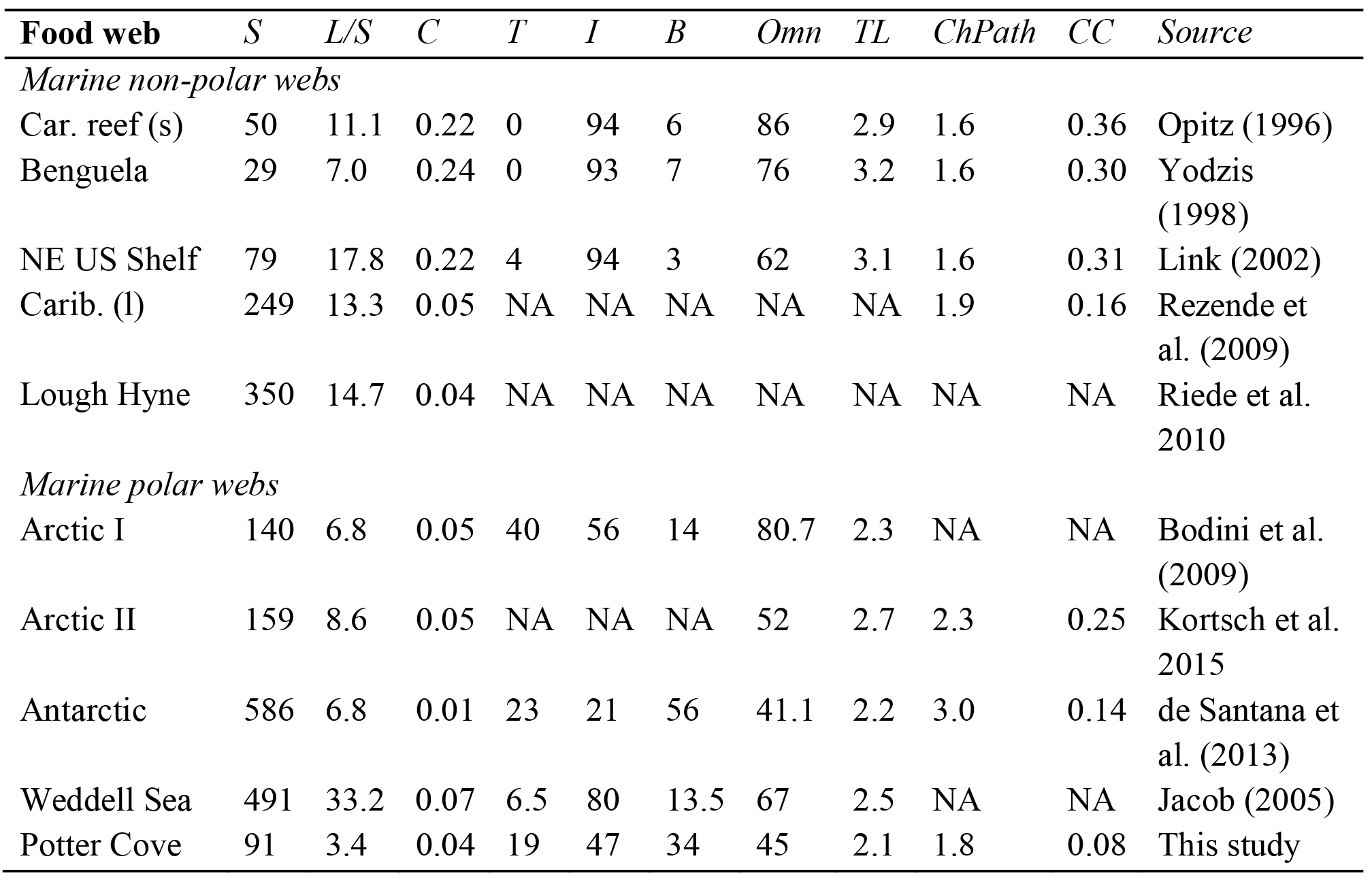
Comparison of network properties between Potter Cove and other marine FWs. S = number of trophic species, L/S = linkage density, C = connectance (L/S^2^), T = % top species, I = % intermediate species, B = % basal species, Omn = percentage of omnivorous, TL = mean trophic level, ChPath = characteristic path length, CC = clustering coefficient. NA: not available data.

| Food web | S | L/S | C | T | I | B | Omn | TL | ChPath | CC | Source |
| --- | --- | --- | --- | --- | --- | --- | --- | --- | --- | --- | --- |
| *Marine non-polar webs* |  |  |  |  |  |  |  |  |  |  |  |
| Car. reef (s) | 50 | 11.1 | 0.22 | 0 | 94 | 6 | 86 | 2.9 | 1.6 | 0.36 | Opitz (1996) |
| Benguela | 29 | 7.0 | 0.24 | 0 | 93 | 7 | 76 | 3.2 | 1.6 | 0.30 | Yodzis (1998) |
| NE US Shelf | 79 | 17.8 | 0.22 | 4 | 94 | 3 | 62 | 3.1 | 1.6 | 0.31 | Link (2002) |
| Carib. (l) | 249 | 13.3 | 0.05 | NA | NA | NA | NA | NA | 1.9 | 0.16 | Rezende et al. (2009) |
| Lough Hyne | 350 | 14.7 | 0.04 | NA | NA | NA | NA | NA | NA | NA | Riede et al. 2010 |
| *Marine polar webs* |  |  |  |  |  |  |  |  |  |  |  |
| Arctic I | 140 | 6.8 | 0.05 | 40 | 56 | 14 | 80.7 | 2.3 | NA | NA | Bodini et al. (2009) |
| Arctic II | 159 | 8.6 | 0.05 | NA | NA | NA | 52 | 2.7 | 2.3 | 0.25 | Kortsch et al. 2015 |
| Antarctic | 586 | 6.8 | 0.01 | 23 | 21 | 56 | 41.1 | 2.2 | 3.0 | 0.14 | de Santana et al. (2013) |
| Weddell Sea | 491 | 33.2 | 0.07 | 6.5 | 80 | 13.5 | 67 | 2.5 | NA | NA | Jacob (2005) |
| Potter Cove | 91 | 3.4 | 0.04 | 19 | 47 | 34 | 45 | 2.1 | 1.8 | 0.08 | This study |

## 4. Discussion

### 4.1. Food web complexity and structure

Potter Cove FW properties of complexity and structure showed several singularities that make the web unique in terms of species-richness, link configuration and topological characteristics. Network complexity was mainly assessed by linkage density (L/S) and connectance (C). Both of these properties were found to be relatively low in the Potter Cove web: L/S=3.4 and C=0.04. Nevertheless, direct comparisons of linkage density and connectance values suggest that marine FWs tend to resemble each other, and that they are fundamentally different from other kinds of FWs, based on their high values (Dunne et al. 2004). Opposite to this hypothesis of marine FW similarity, de Santana et al. (2013) found that connectance in the Arctic marine FW was 5 times larger than that of the Antarctic one (0.05 versus 0.01). Furthermore, within marine webs, polar networks tend to display low values of linkage density (de Santana et al. 2013). In this sense, complexity exhibited in the Potter Cove FW resembles closely to what is known so for FWs in Polar regions.

Could low values of linkage density and connectance in Potter Cove network be a consequence of methodological issues? Dunne et al. (2002b) suggested that both low- and high-connectance FWs are unusual, and that extreme connectances may sometimes be artifacts of assembly procedures. They exemplified this statement by showing that the lowest connectance webs they studied (C ≈ 0.03, Grassland and Scotch Broom), which is similar to Potter Cove FW connectance value, are “source-webs”. These are constructed by linking food chains upward starting from one or a few basal species. The Potter Cove FW is a species-rich ecological network and not a source-web since it was not constructed upward from one or two basal species but it is characterized by > 30% basal species. Thus there is no evidence we know of which suggests that our low values of linkage density and connectance were a consequence of the assembly procedure of the network. In turn this implies that the assembly-connectance relationship in FWs is not as strong as previously thought (Dunne et al. 2002b).

Whether ecological networks display low or high L/S and C values is crucial to gain insight in the ecosystem’s synthetic properties like robustness. Empirical analyses of FWs support the notion that the robustness of a FW increases with its linkage density and connectance (De Angelis 1975, Dunne et al. 2002a, Montoya and Solé 2003). Low values of L/S and C found in Potter Cove FW, combined with ongoing climate change effects on benthic communities in the area (Pasotti et al. 2015b, Sahade et al. 2015), suggest potential ecosystem fragility which need to be addressed.

Furthermore, direct comparison of common FW properties, like percentages of top, intermediate and basal species, indicates that the Potter Cove network has strong structural differences and shows unique features compared to other marine ecosystems. Important dissimilarities were found in top and basal species values as Potter Cove FW shows a higher number of these trophic species. After comparing 19 FW properties, Dunne et al. (2004) concluded that the excessively low percentage of basal taxa in marine FWs compared to other systems is clearly an artifact of poor resolution of primary producers and consumer links to them. One of the methodological strengths of Potter Cove FW is the high taxonomic resolution of the basal nodes. A good taxonomic resolution of the lower trophic levels, such as the macroalgal community, is essential to understand Potter Cove ecosystem functioning, since there seems to be a species-specific selective consumption (Barrera-Oro and Casaux 1990, Iken et al. 1997, Iken et al. 1998). Furthermore, algal species show a marked pattern of depth distribution and tridimensional structure (Quartino et al. 2005, Huang et al. 2007). Macroalgae are one of the main primary producers in Potter Cove, and probably support a large fraction of secondary production of the benthos community (Quartino et al. 2008). Implications in ecosystem functioning and stability are only possible to elucidate in FWs where species involved in energy and matter transfer processes are well represented.

Proportions of intermediate species (I) and omnivory (*Omn*) in Potter Cove FW are relatively low when compared to other marine webs, but close to values for Antarctic FW as reported by de Santana et al. (2013). Levels of *I* and omnivory are usually correlated in FW studies, as the majority of species acting as predators and prey also feed on more than one trophic level (omnivorous). The importance of omnivory for the structure and dynamics of FWs is a long-standing controversy in ecology (Burns 1989, Polis 1991), and whether omnivory stabilizes or destabilizes webs is not clear (Vandermeer 2006, Namba et al. 2008, Johnson et al. 2014). In Antarctica a recent study suggests that omnivory is a beneficial trait as it allows for more responsive and flexible utilization of food sources that may be temporally and spatially constrained and unpredictable (Norkko et al. 2007). The omnivory reported here for Potter Cove FW is the second lowest percentage among marine webs included in the present study, would suggest a low stability for Potter Cove FW. Additionally, this result generates testable hypotheses about the probable stabilizing role of omnivory in large communities, since it was proven that the risk of secondary extinctions after primary loss of species depends on the trophic position of the extinct species (Borrvall et al. 2000) and the diversity of that trophic level (insurance hypothesis, Yachi and Loreau 1999).

The mean trophic level for this FW (2.1) is also relatively low, which is the result of several singularities of the Potter Cove ecological network. Firstly, as already mentioned, the number of basal trophic species is high, exceeding 30% of number of species (diversity). What´s more, the maximum trophic level was 4.27, lower than most other FWs studied (Dunne et al. 2002b, 2004), which implies that top and basal species are separated by few intermediate taxa. It is worthy to clarify here that Antarctic top predators, e.g. marine mammals, might increase maximum trophic level of the web but were not included as they are rarely reported in the Cove. Therefore, the transfers of energy or nutrients from the base to the top of Potter Cove FW is small, so that the number of times chemical energy is transformed from a consumer's diet into a consumer's biomass along the FW is also small. Another reason why the mean trophic level is low is the fact that most predators at intermediate levels (e.g. amphipods, isopods, bivalves, *N. coriiceps*) feed predominantly on algae species and/or detritus, being mainly the product of dead and decomposed macroalgae in Potter Cove (Iken et al. 1998, Huang et al. 2006, Quartino et al. 2008). The macroalgal detritus decomposes and is eaten by detritivores and suspensivores (e.g. sponges, ascidians, bryozoans, cnidarians), supporting an important amount of the secondary production (Tatián et al. 2004). The obtained low mean trophic level for Potter Cove FW clearly shows what species-specific and/or community studies have suggested. These characteristics of ecological communities have a high impact on ecosystem functioning, such as nutrient and carbon cycling, and trophic cascades (Post 2002).

Short characteristic path length for Potter Cove FW (≈ two degrees of separation) is similar to lengths found in other marine FWs. The length between pairs of species within marine webs is low (≈1.6 links) compared to other types of FWs, with values ranging from 1.3 to 3.7 (Dunne et al. 2002b). This suggests that most species in Potter Cove FW are potentially very close neighbours, and that negative effects could spread rapidly and widely throughout the web (Dunne et al. 2002a).

Additionally, the clustering coefficient in this web (0.08) was an order of magnitude lower than those reported for other marine FWs (Link 2002, Dunne et al. 2004). A low coefficient indicates that most species are similarly connected to each other, i.e. there are no densely sub-groups of species interacting with one other. Particularly, the clustering coefficient result of Potter Cove FW might be the consequence of hubs (i.e. species with high degree, > 20 links) connected with most of the species across the web and not with a specific group of species. The most connected species, *N. coriiceps* (demersal fish) and *Ophionotus victoriae* (brittle star), have the widest ecological niches in our study, being generalists and omnivores. By feeding across several trophic levels and transversely in the FW, these species have a strong effect on clustering. Specifically, *N. coriiceps probably* represents a keystone species in the bentho-pelagic coupling process promoting the transfer of matter and energy between habitats (Barrera-Oro and Casaux 2008). At the same time, these hub species might be essential for understanding the spread of perturbations (i.e. biodiversity loss) through the entire FW network.

### 4.2. Degree distribution and overlap graphs: implications for ecosystem functioning

Webs with low connectance (*C* ≈ 0.03), such as Potter Cove FW, are more likely to display power law degree distributions (Dunnet et al. 2002a, Montoya and Solé 2002), consistent with the small-world phenomenon. These are webs combining high clustering, like regular lattices and short path length, like random graphs (Watts and Strogatz 1998). Therefore, the Potter Cove FW, with a low estimated connectance (*C* = 0.04), should display a power law degree distribution. However, it fits best to an exponential distribution according to the low clustering coefficient. The existence of a universal functional form in the degree distribution of FWs is still under debate, though Stouffer et al. (2005) have shown that approximately exponential degree distributions can be derived from two different models: nested-hierarchy and generalized cascade.

The influence of the degree distribution on the vulnerability of complex networks against random failures and intentional attacks has become well known since the work of Albert et al. (2000). Considering this relationship between degree distribution and vulnerability, Potter Cove FW would be highly fragile to the removal of the most connected species, but not as much as in power law networks (Albert el al. 2000, Dunne et al. 2002a, Estrada 2007). Furthermore, de Santana et al. (2013) suggested that less connected communities should be more sensitive to the loss of basal species than complex communities because the consumers in simple communities are dependent on only a few species and cannot survive their loss. Nevertheless, we hypothesize that although Potter Cove FW shows low connectance, it will be robust against basal node extinctions due to the high percentage of these trophic species.

In addition, degree distribution of links in the competition graph showed that most species have limited diets, feeding exclusively on few prey, whereas few species feed on a large amount of food-sources, usually being generalists. The graph suggests that several predator species have high prey overlap and thus the potential to strongly interact and compete for common prey; this is the case for sponges, demersal fish and amphipods. We focused the analysis on fish and amphipods as they are known to play an important role on the Antarctic marine ecosystem (Barrera-Oro and Casaux 1998, Momo et al. 1998, Barrera-Oro 2002, Huang et al. 2006). Fish data reflects that there is dietary overlap between *N. coriiceps* and *H. antarcticus* on the one hand and between *Trematomus newnesi* and *N. rossii* on the other hand. Most of the dietary comparisons for demersal Antarctic fish communities have dealt with food overlap between fish species pairs (Barrera-Oro 2003). Dietary overlap index (“S” index of Linton et al. 1981) between *N. coriiceps* and *N. rossii* in Potter Cove as estimated by Barrera-Oro (2003) was 55%, meaning that these species could compete for more than half of their food-sources. The same study estimated the index for *N. coriiceps* – *T. newnesi*, being 18%, and *N. coriiceps* – *H. antarcticus,* being 19%. Barrera-Oro (2003) concludes that there is no evidence of food competition among the shallow cold-water fish communities in Potter Cove. Nevertheless, our results show that *N. coriiceps* and *H. antarcticus* have many prey in common, with a high degree of overlapping. However, due to the differences in mobility, habitat use and adult size between these two species (total length: *45 and 13 cm respectively*), competition is probably low (Casaux 1998, Barrera-Oro 2003). Although the first one is a generalist and the latter a specialist, both species can be grouped in the same feeding category given that they are benthos feeders, eating amphipods (e.g. *Gondogeneia antarctica, Paradexamine sp., Prostebbingia sp, Eurymera monticulosa*), gastropods (e.g. *Margarella antarctica, Nacella concinna, Eatoniella sp, Neobuccinum eatoni*), polychaetes (e.g. Nereidae), and krill in summer (*Euphausia superba*). On the other hand, the competition graph for amphipods exhibited low dietary overlap among species. It is worth mentioning that hyperiids and *Bovallia gigantea* are not connected, which indicates that they do not share food sources with any other amphipods, nor between themselves. Hyperiids and *B. gigantea* are both carnivores, though the latter mainly feeds on other species of amphipods, such as *E. monticulosa*, *Prostebbingia sp.* and *G. antarctica* (Richard 1977). On the contrary, hyperiids principally eat planktonic prey, such as copepods (Pakhomov and Perissinotto 1996). The most important result of the overlap graph is that species are separated according to their feeding strategies: herbivores (*P. gracilis, G. antarctica, O. bidentata* and *Prostebbingia sp.* – left of the graph), detritivores (*C. femoratus* and *Paradexamine sp.* – middle graph), and scavengers (*W. obesa*, *H kergueleni*, *O. plebs* and *P. integricauda* – right of the graph). This demonstrates the importance and utility of the analysis of competition graphs, in order to better understand alternative energy pathways within apparent trophic guilds; analysis that would be improved by adding information on each predator species (e.g. body size and mass, niche specialization).

Common-enemy graph derived from Potter Cove FW showed a hyper-connected graph, which implies that most prey species share at least one predator. The fact that the prey overlap graph of this FW exhibited high connectivity and exponential distribution has implications for the functioning of the ecosystem. High-connected prey in Potter Cove FW are: phytoplankton – zooplankton, benthic diatoms – epiphytic diatoms, and fresh detritus – benthic diatoms. The latter shows that several sources of food and alternative energy pathways exist in the Potter Cove ecosystem: phytoplankton (Ahn et al. 1993), benthic microalgal production (Dayton et al. 1986, Gilbert 1991), and horizontal advection of allochtonous food particles (Dunbar et al. 1989); important sources of organic matter for marine organisms living in coastal Antarctic ecosystems.

In conclusion, comparison of FW properties revealed a particular combination of characteristics for the Potter Cove ecological network: middle size (*S* ≈ 100) compared to other marine FWs, low linkage density and connectance (with no evidence of being an artifact of resolution or assembly procedure), low %-omnivory, short path length and low clustering coefficient. According to the overlap graphs and their degree distributions, and the consistency with field observations and investigations, we suggest these analyses are useful tools to gain insight into ecosystem functioning. What is more interesting, the common-enemy graph showed the existence of alternative energy pathways consistent with field investigations in the Cove. As also suggested for East Antarctica FW (Gillies et al. 2012), carbon flow among the benthic fauna in Potter Cove is complex, with multiple sources of carbon being utilized, which can be asserted given the good basal resolution of our network.

From a network perspective, Potter Cove FW properties suggest fragility and potential trophic cascade effects although multiple energy pathways might add robustness to the web. Our results suggest that species with a high number of links (e.g. *Notothenia corriceps, Ophionotus victoriae, Gondogeneia antarctica*) could be considered as keystone species for the robustness of Potter Cove ecosystem.

## Acknowledgments

This research was supported by Consejo Nacional de Investigaciones Científicas y Técnicas (CONICET, Argentina), Universidad Nacional de General Sarmiento and Alfred Wegener Institute for Polar and Marine Research (AWI, Germany). The work was partially funded by PIO 14420140100035CO CONICET Argentina and conducted in the frames of the EU research network IMCONet funded by the Marie Curie Action IRSES (FP7 IRSES, Action No. 319718). We thank Dave K.A. Barnes for constructive suggestions on language aspects, which helped us to improve the manuscript.

## Appendices

Supplementary material Appendix A and B are available at DOI 10.6084/m9.figshare.4498715.

